# Insulin signalling activates multiple feedback loops to elicit hunger-induced feeding in Drosophila

**DOI:** 10.1101/364554

**Authors:** Sreesha R. Sudhakar, Jishy Varghese

## Abstract

Insulin, a highly conserved peptide hormone, links nutrient availability to metabolism and growth in animals. Besides this, in fed states insulin levels are high and insulin acts as a satiety hormone. In animals that are food deprived insulin levels remain low which facilitates hunger induced feeding. Contrary to expectations, we present evidence for persistent *Drosophila insulin*-*like peptide* gene expression and insulin signalling during initial phases of starvation. Maintenance of insulin signalling is crucial to sustain feeding responses during initial stages of starvation. Insulin signalling acts in a feedback loop involving the abdominal fatbody to maintain *dilp* gene expression in the early stages of food deprivation. Furthermore, another feedback regulatory loop between insulin-producing cells (IPCs) and neurons that produce the orectic hormone short-neuropeptide-F (sNPF), maintains sNPF levels and triggers feeding behavior. Thus, insulin acts through multiple feedback regulatory loops to elicit orexigenic responses and aid in efficient utilization of energy stores during early starvation.

## Introduction

Energy is indispensable for all biological functions and the central metabolic pathways that aid in cellular energy production is highly conserved in all organisms. Animals have to cope up with the fluctuating availability of food resources, which requires precise sensing of internal energy levels and efficient regulation of nutrient uptake. A negative energy balance would promote nutrient intake and restrict energy expenditure, and vice versa (1). Hunger, an adaptive response to food deprivation, triggers various stereotypic behaviours to meet the nutritional and energy demands of higher animals. Shortages in the supply of food would enhance foraging, trigger an urge to feed and motivate animals to consume noxious foods which are normally avoided (2). Besides this, internal energy reserves aids in meeting the physiological needs of an organism during starvation. A plethora of complex communication networks that include various neuropeptides and hormones play pivotal roles in eliciting behavioural responses and manage energy metabolism during starvation (3-8).

Insulin, a major peptide hormone, aids in energy sensing and maintains nutrient homeostasis by controlling a myriad of cellular, metabolic and physiological processes. Insulin is highly conserved across metazoans, including the widely used model organism Drosophila (9-12). In Drosophila, a set of 14 neurosecretory cells, present at the *pars intercerebralis* region of the brain, produce Drosophila insulin-like peptides - DILPs (13, 14). Drosophila genome contains eight *dilp* genes that encode DILPs1-8, of which *dilp2*, *dilp3* and *dilp5* are expressed mainly by the neuronal IPCs (15). Drosophila IPCs function as a central node in maintaining nutrient homeostasis as various nutrient signals originating from the fatbody, an endocrine organ similar to vertebrate liver and adipocytes, targets these cells (16). Based on the fatbody derived nutrient signals IPCs manage several cellular and physiological functions like growth, metabolism, stress responses, feeding, etc., (17-19). Furthermore, an appetite-stimulating neuropeptide, sNPF, orthologous to mammalian Neuropeptide-Y (NPY) controls IPCs and promotes growth by triggering *dilp* gene expression (20). However, investigations on the role of Drosophila IPCs in sNPF mediated feeding control are only beginning to emerge (21).

Here, we unravel a novel role for insulin in hunger-induced feeding responses. Insulin is believed to act as a ‘satiety hormone’ and insulin release induced by feeding marks a high-energy state in various animals including Drosophila (22-24). Contrary to this, we show that insulin signalling remains high during initial phases of starvation in adult Drosophila, which is crucial for hunger induced food intake. Furthermore, during early stages of starvation insulin signalling acts on the abdominal fatbody to maintain DILP levels and enhance the drive to feed. The levels of DILPs and insulin signalling that stimulate feeding during early stages of nutrient deprivation is maintained by the action of neuropeptide sNPF on IPCs. Together with this, insulin signalling also maintains the expression of ‘orexigenic’ neuropeptide sNPF. Thus, insulin acts as a ‘hunger hormone’ during initial stages of food withdrawal and multiple feedback loops activated by insulin signalling maintains feeding during this stage.

## Results

### Insulin signalling remains high during early phases of starvation

Hunger enhances the drive to feed. As expected, there was enhanced feeding intake in flies (5-day old males) reintroduced to food after nutrient deprived for 12, 24 and 36 hours (Fig 1). Previously reported findings show that DILPs produced by IPCs regulate feeding in Drosophila and DILPs may act as an anorexic hormone (23, 25). Hence, we tested *dilp2*, *dilp3 and dilp5* mRNA levels during starvation, as they were shown to be nutrient sensitive by earlier reports (13, 25-28). In contrast to our expectations, we found no decrease in the levels of *dilp* mRNAs after 12 hours of starvation in comparison to fed flies (Fig 2A). In fact, we observed a significant increase in the transcript levels of *dilp3* after 12 hours of starvation. Even after 24 and 36 hours of starvation we saw a significant reduction of only *dilp5* mRNA levels (Fig 2A). To measure DILP protein levels in the IPCs, we used a peptide antibody against DILP2 during starvation. DILP2 protein levels accumulate in the IPCs leading to a reduction in their levels in the hemolymph in response to starvation (17). However, we observed no changes in DILP2 protein levels in the IPCs after 12 hours of starvation (Fig 2B), suggesting that at this stage there is no effect on the release of DILP2 protein into the hemolymph. Thus, flies starved for 12 showed hunger-driven feeding (Fig 1), despite the fact that *dilp* gene expression and the levels of DILP2 protein remained unaltered in the IPCs during this stage (Fig 2A and B). This prompted us to check the activity of insulin signalling pathway during early stages of starvation.

**Fig. 1.**
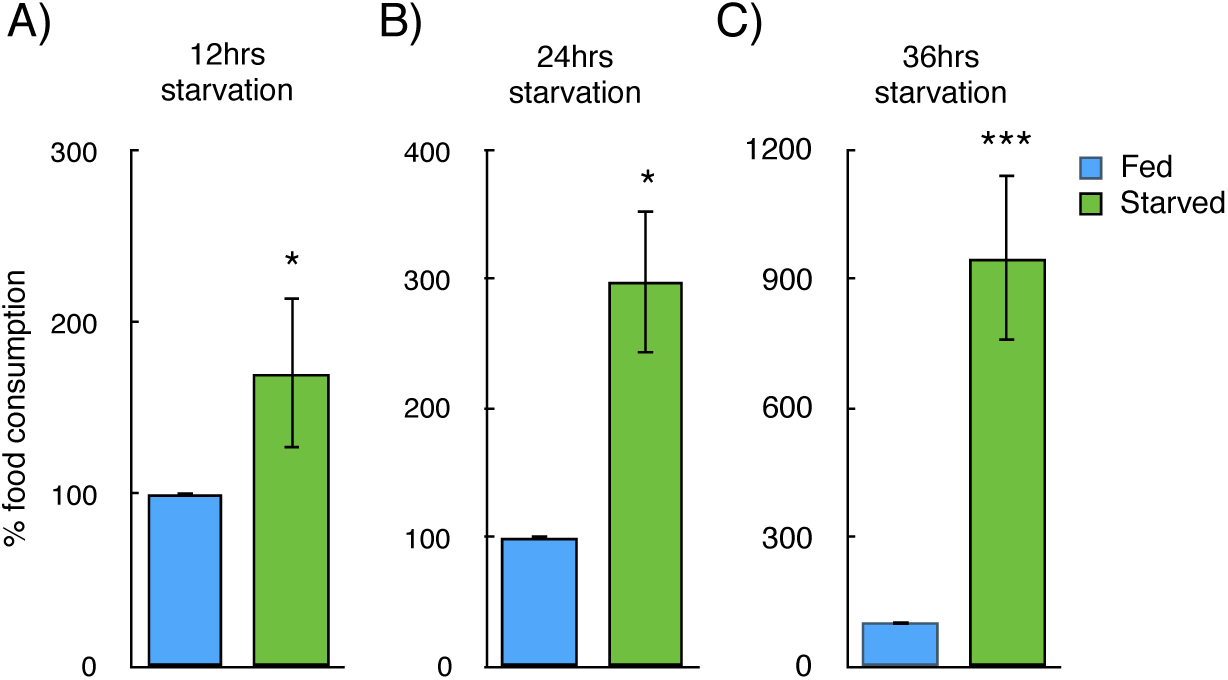
Food withdrawal triggers hunger induced feeding responses in adult Drosophila. (**A**) Enhanced food consumption by flies food deprived for 12 hours [n>20], (**B**) 24 hours [n=8] and (**C**) 36 hours were reintroduced to food [n>20], values are in percentage normalized to flies that are fed on normal food. (P-value ^∗^<0.05; ^∗∗^ <0.01,^∗∗∗^ <0.005, n=number of biological replicates; error bars represents SEM)

**Fig. 2.**
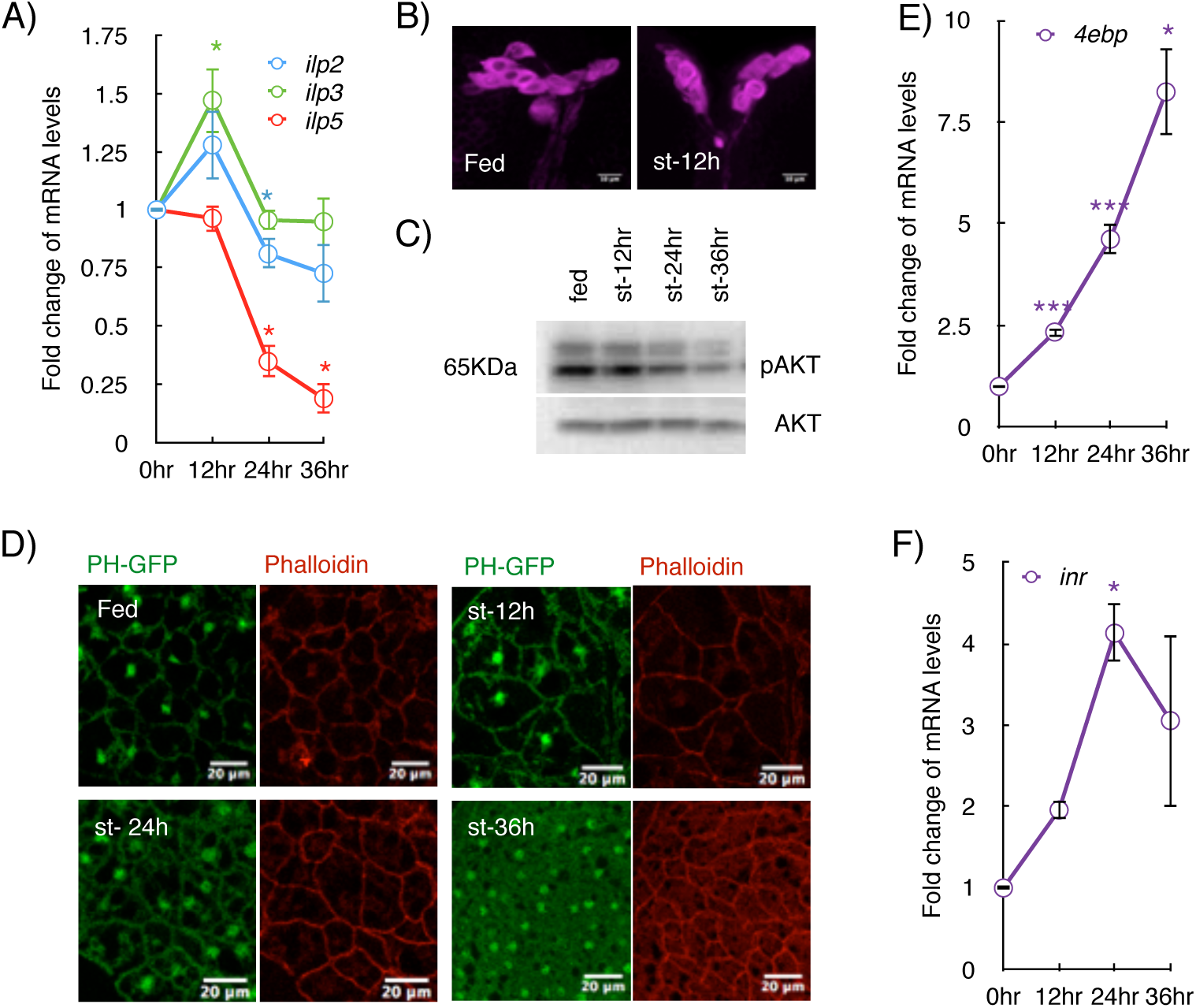
Insulin gene expression and activity of insulin signalling pathway remains high during initial stages of food deprivation. (**A**) *dilp2*, *3 and 5* mRNA levels during starvation, values show fold change in transcript levels after starving flies for various time duration mentioned (in hours). Only *dilp5* mRNA levels show significant reduction after 24 hours of starvation. [n=6] (**B**) DILP2 protein levels in the IPCs in fed and 12 hours starved flies. [n=7] (**C**) Western blot showing phospho-AKT levels in lysates obtained from fed and starved flies (after indicated durations of starvation), total AKT is shown as a control for protein levels. Phosphorylated levels of AKT do not change after 12 hours of starvation. [n=3] (**D**) PH-GFP (Green) localization in the abdominal fat body of flies that were fed normally or starved for 12, 24 and 36 hours, the fatbody cells were stained with Phalloidin-Alexa568 (red) [n>8] (**E**) *4ebp* and (**F**) *inr* mRNA levels during indicated hours of starvation [n>7], values show fold change in transcript levels, compared to non-starved flies (0 hours). (P-value ^∗^<0.05; ^∗∗^ <0.01,^∗∗∗^ <0.005, n=number of biological replicates; error bars represents SEM)

To measure the levels of insulin signalling in response to starvation, we used multiple strategies. We assayed AKT phosphorylation in adult flies as a readout of insulin signalling using an antibody specific to phospho-AKT. Insulin signalling leads to phosphorylation and activation of AKT via phosphatidylinositol-3 kinase (PI3K). We observed minimal changes to AKT phosphorylation (Fig 2C) during early stages of starvation (12 hours), a stage where flies show an enhanced urge to feed. However, in response to long periods of food deprivation (24 or 36 hours) phospho-AKT levels showed a substantial decrease, whereas, total AKT levels remained unaltered (Fig 2C). Further, to monitor the levels of insulin signalling during starvation, we checked the localization of PH-GFP (pleckstrin-homology domain-GFP, an indicator of insulin/PI3K activity) in the adult fatbody (29). In cells that receive high levels of insulin PH-GFP would localize to the plasma membrane and upon reduction of insulin signalling PH-GFP would show a decrease in membrane localization. We observed only minimal changes to PH-GFP membrane localization after 12 hours of starvation in adult fatbody (Fig 2D). In comparison, the membrane localization of PH-GFP was disrupted after 24 hours of starvation (Fig 2D).

Further, to confirm the levels of insulin signalling during early phases of starvation, we checked the expression of two insulin target genes, *insulin receptor (inr)* and *4E binding protein (4ebp)*, known to be repressed by insulin signalling (30, 31). We found that *4ebp* and *inr* mRNA levels increased moderately after 12 hours of starvation (Figure 2E and F), confirming that insulin signalling underwent only modest changes during early stages of starvation. The levels of *4ebp* and *inr* mRNA increased substantially only after 24 hours of starvation, indicating a sharp reduction in insulin signalling during late stages of starvation.

As an anorectic hormone, insulin levels and insulin signalling were expected to drop in nutrient-deprived animals (2, 23, 25). However, we observed only moderate changes to insulin levels and insulin signalling during early stages of starvation, even though at this stage the flies exhibited enhanced feeding responses. The enhanced feeding and simultaneous retention of insulin signalling activity suggested that insulin may aid in feeding during early phases of starvation. To address this, we tested if insulin signalling promotes hunger-induced feeding during early starvation.

### Insulin-Producing Cells aid in enhanced feeding in response to starvation

To analyze the role of insulin signalling in hunger-induced feeding insulin activity was blocked in flies subjected to food deprivation. We used a tubulin-promoter driven temperature-sensitive allele of Gal80 (*tub*-*Gal80^ts^*) and temporally restricted the expression of apoptotic gene *reaper* to adult IPCs to minimise developmental or growth-related effects (for details, please see S1 Fig). In the following experiments, we used *tub*-*Gal80^ts^* to restrict the expression of various transgenes to adult flies to avoid developmental effects, unless mentioned otherwise. Expression of *reaper* in the IPCs just prior to starvation, resulted in a significant reduction of hunger-induced food consumption in flies starved for 12 hours (Fig 3A). The inactivation of IPCs was confirmed by measuring gene expression of *dilp2*, *dilp3*, *dilp5* and insulin target gene *4ebp* (data not shown). Also, the expression of *Kir2.1*, a hyperpolarizing potassium ion channel which blocks neuronal activity, in the IPCs hindered hunger-induced feeding in response to 12 hours of food deprivation (Fig 3A).

**Fig. 3.**
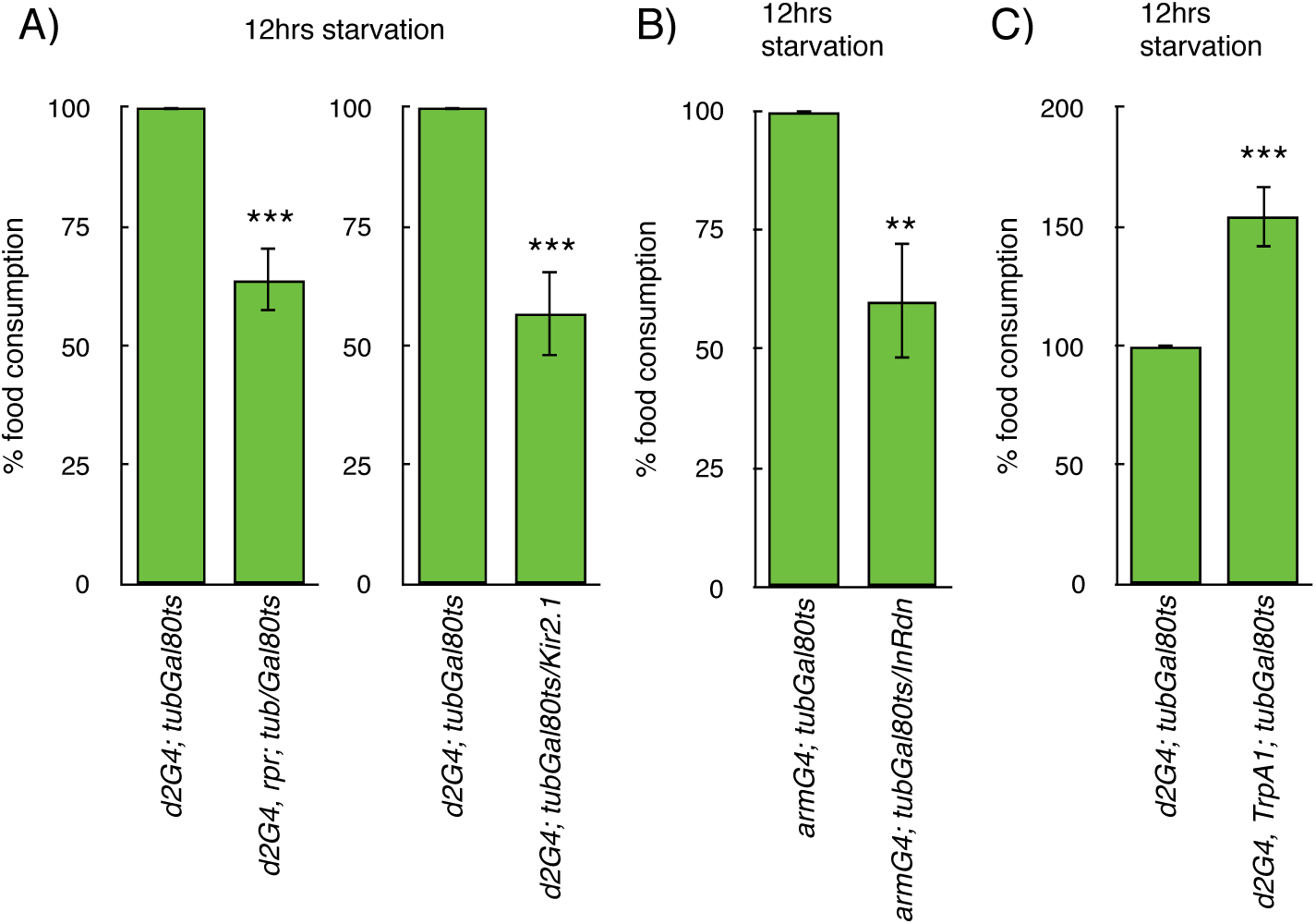
IPC function and insulin signalling during early phases of starvation is inevitable for hunger induced feeding. (**A**) blocking IPC function by expressing *reaper* or *Kir2.1* inhibited feeding induced by 12 hours of starvation [n=13 and 6] (**B**) inhibiting insulin signalling in the whole body by ubiquitous expression of an insulin receptor dominant negative transgene (*inr*-*DN*) decreased feeding induced in response to 12 hours of starvation [n=11] (**C**) *TrpA1* mediated activation of IPCs led to enhancement of feeding induced by 12 hours of starvation [n=15] (*All values are in percentage normalised to control for starved conditions*, *n*=*number of biological replicates P*-*value* ^∗∗^<*0.01*; ^∗∗∗^ <*0.005*; *error bars represents SEM)*

Further, we blocked InR function/expression in the whole fly to limit systemic insulin-signalling during starvation. We expressed UAS-*inr*-RNAi and UAS-InR-dominant negative (*inr*-*dn*) transgenes precisely during the food withdrawal stages using a ubiquitous *armadillo* Gal4 driver under the control of *tub*-*Gal80^ts^* (Fig 3B, S2 Fig A). A temperature sensitive transheteroallelic *inr* mutant, *inr^E19/GC25^* was used to assess the effects of inactivating InR function (S2 Fig A). Switching *inr^E19/GC25^* flies from 18°C to 29°C renders *inr* gene inactive, and we shifted the temperature of the adult flies just prior to the starvation period to inactivate *inr* function (32). All the above approaches which blocked insulin signalling systemically led to a decrease in hunger-induced feeding in response to 12 hours of food withdrawal (Fig 3B and S2 Fig A). Thus, our results show that IPC function and maintenance of insulin signalling are crucial for hunger-induced feeding at the initial phases of starvation. Based on these observations, next, we addressed whether insulin acts as an orexigenic hormone during early starvation.

### Drosophila insulin-like peptides triggers feeding

Earlier reports show that enhancing DILP levels resulted in anorexigenic effects in larvae and flies (23, 25). In our experiments overexpression of the cationic channel - TrpA1 (transient receptor potential A1), a neuronal activator, in the IPCs led to an increase in food consumption in flies that were food deprived for 12 hours (Fig 3C). Overexpression of *dilp2* in adult IPCs also increased feeding in flies that were nutrient deprived for 12 hours (S2 Fig B). However, overexpression of DILP2 reduced feeding in flies starved for 36 hours (S2 Fig B), in agreement with the previous studies (23, 25). Overall these results provide evidence that DILPs promote feeding in flies only during early stages of starvation. Whereas in later stages of food deprivation insulin blocked hunger-induced feeding as shown previously (23, 25). Also, when insulin was overexpressed throughout developmental phases, there was a reduction in hunger-induced feeding (data not shown), similar to earlier reports (23). Given such crucial roles for insulin in feeding we addressed the mechanisms by which insulin triggers feeding responses during starvation.

### IPCs and abdominal fatbody acts in a positive feed-back loop to trigger feeding and manage *dilp* gene expression

To explore the means by which insulin aids in hunger-induced feeding, we blocked insulin signalling in various adult tissues. We targeted insulin signalling in the IPCs, head and abdominal fatbody, tissues that aid in maintaining nutrient homeostasis in adult flies. As mentioned earlier, limiting InR levels in the whole fly led to a reduction in starvation-induced feeding (Fig 3B and S2 Fig A). Blocking InR in the abdominal fatbody led to the reduction of starvation triggered feeding (Fig 4A), whereas, no effects on feeding was observed by blocking InR in the head fatbody or IPCs (S4 Fig A and B). For details of temporal restriction of the expression of abdominal and head fatbody drivers, please see S3 Fig. Also, blocking insulin signalling by expressing PTEN in the abdominal fatbody led to a reduction of starvation triggered feeding (Fig 4A). Thus, reducing insulin signalling in the abdominal fatbody in flies nutrient deprived for 12 hours negatively affected feeding responses. Food deprivation for 12 hours of starvation did not lead to a reduction in *dilp* mRNA levels (Fig 2A). Hence, we tested if insulin signalling in the abdominal fatbody during the early phases of starvation aids in the maintenance of *dilp* mRNA levels. Blocking InR in the adult abdominal fatbody led to a reduction of *dilp2*, *dilp3* and *dilp5* mRNA levels in flies starved for 12 hours (Figure 4B). Thus, insulin signalling in the abdominal fatbody in flies nutrient deprived for 12 hours is crucial for feeding responses and maintenance of *dilp* mRNA levels. Our data show that IPCs and abdominal fatbody acts in a feedback loop to maintain DILP levels. Next, we decided to explore the mechanisms by which the adult fatbody would regulate feeding during early stages of starvation.

**Fig. 4.**
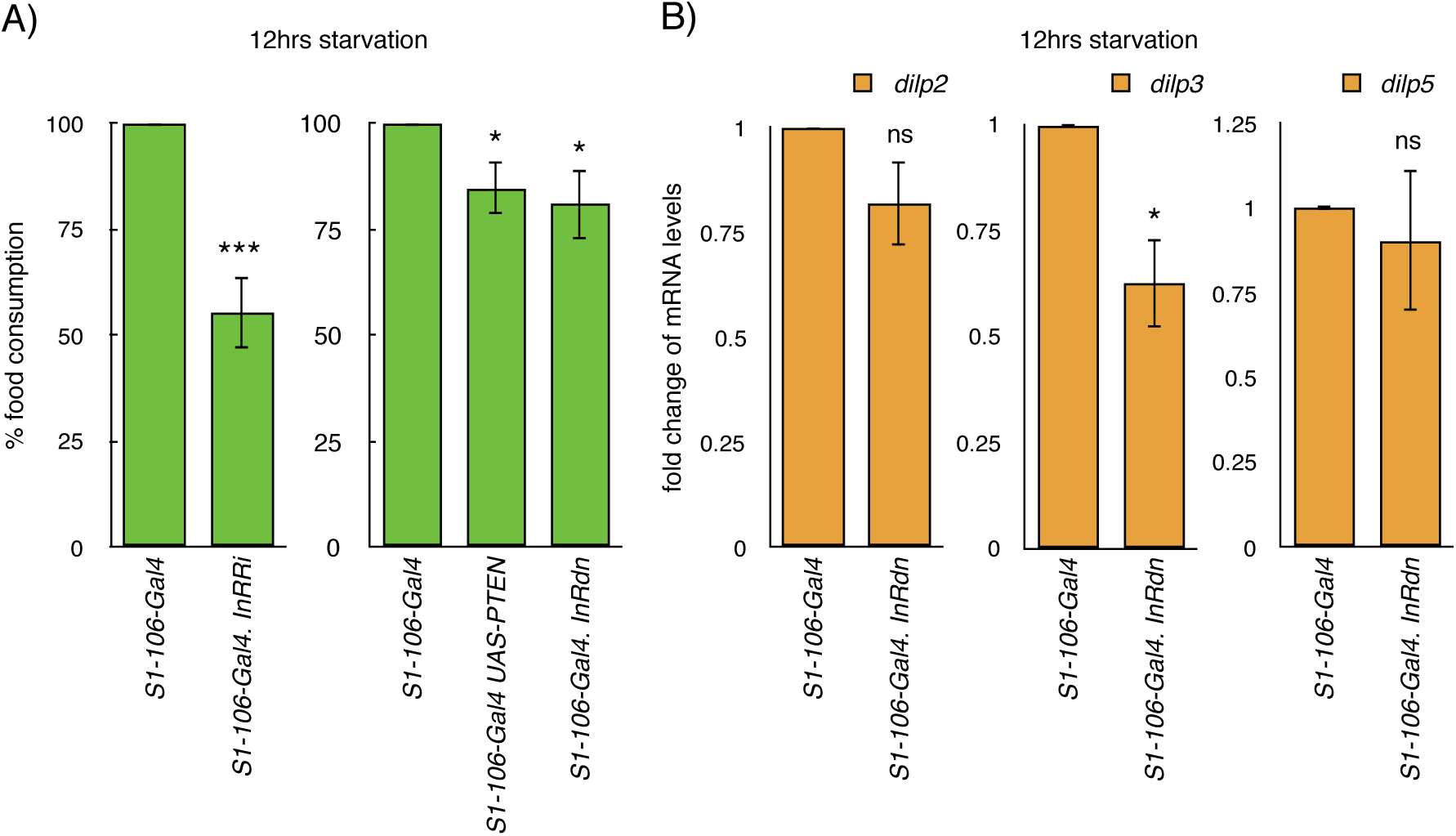
Mechanisms of insulin action that aid in hunger induced feeding. (**A**) Food consumption in response to 12 hours of starvation was reduced when *inr*-*RNAi*, *inr*-*dn* and *UAS*-*PTEN* were expressed in the abdominal fat body using *S_1_106*-*Gal4* [n=10 and >30] and (**B**) After 12 hours of starvation *dilp2*, *dilp3* and *dilp5* mRNA levels also showed a decrease in response to blocking insulin signalling in the abdominal fat body, values are normalized to control genotypes [n=5]. (*All values are in percentage normalised to control or heterozygous genetic background for starved conditions*, *P*-*value* ^∗^<*0.05*; ^∗∗^<*0.01*; ^∗∗∗^ <*0.005*; *n*= *number of biological replicates*, *error bars represents SEM).*

Fatbody in Drosophila can sense changes in the nutrient levels and control IPC functions through various humoral signals (17). In response to changes in nutrient status Drosophila fatbody release metabolic signals like DILP6, leptin homolog Unpaired2 (Upd2), peptide hormone CCHa2 (CCHamide-2) and a TNF-like cytokine Eiger. These fatbody derived signals control the function of IPC and regulate DILP levels (18, 19, 33, 34). To address the role of fatbody derived signals on feeding, we blocked the expression of genes that encode these humoral signals. Blocking *dilp6*, *eiger* or *upd2* mRNA levels using RNAi lines and dominant negative transgenes (whereever they were available) in the abdominal fatbody did not lead to consistent changes to feeding responses in flies starved for 12 hours (S5 Figure). Moreover, overexpression of *dilp6* in the abdominal fatbody did not show any effects on starvation-induced feeding at 12 hours (S5 Figure). Even though our results unravel a role for abdominal fatbody in the regulation of hunger-induced feeding responses, a possible fatbody derived humoral signal (FDS) that act on IPCs to regulate hunger-induced feeding responses remains elusive. Next, we assessed the signalling mechanisms that activate IPCs and enhance feeding in response to starvation.

### A second positive feedback mechanism which triggers feeding during starvation

Drosophila neuropeptide sNPF triggers food intake in larvae and adult flies and is thought to be crucial for hunger driven food intake (25, 35). The receptor for sNPF (sNPFR) is expressed in the IPCs and sNPF signalling in larval IPCs triggers *dilp1 and 2* expression, which results in larval growth (20). The aforementioned facts prompted us to ask if the maintenance of DILP levels during early stages of starvation is due to high sNPF signalling. We addressed this using multiple approaches, starting with blocking sNPF signalling in the IPCs of flies starved for 12 hours, which led to a reduction of hunger-induced feeding (Fig 5A). However, in contrast to early stages of starvation hindering sNPF signalling in IPCs did not reduce hunger-induced feeding at 36 hours of starvation (S6 Fig A). Next, we measured *dilp* mRNA levels and insulin signalling, after blocking sNPF signalling in the IPCs in flies which were starved for 12 hours. As an outcome of blocking sNPF signalling in the IPCs the levels of *dilp3* mRNA and insulin signalling were reduced, (Fig 5B), however the effects of blocking sNPF signalling in IPCs on *dilp2 and dilp5* mRNA levels were not significant (Fig 5B). Finally, we asked if sNPF signalling in IPCs maintain *dilp* mRNA levels and thereby enhance feeding in response to starvation. To address this, we increased DILP2 levels when sNPF signalling was blocked in the IPCs, the expression of DILP2 has been shown to be sufficient to rescue the effect of ablation of IPCs (14). Enhancing DILP2 levels during initial hours of starvation was sufficient to limit the effects of blocking sNPF signalling in the IPCs (Fig 5C). From these experiments, we conclude that sNPF signalling in IPCs during early stages of starvation maintains DILP levels and feeding.

**Fig. 5.**
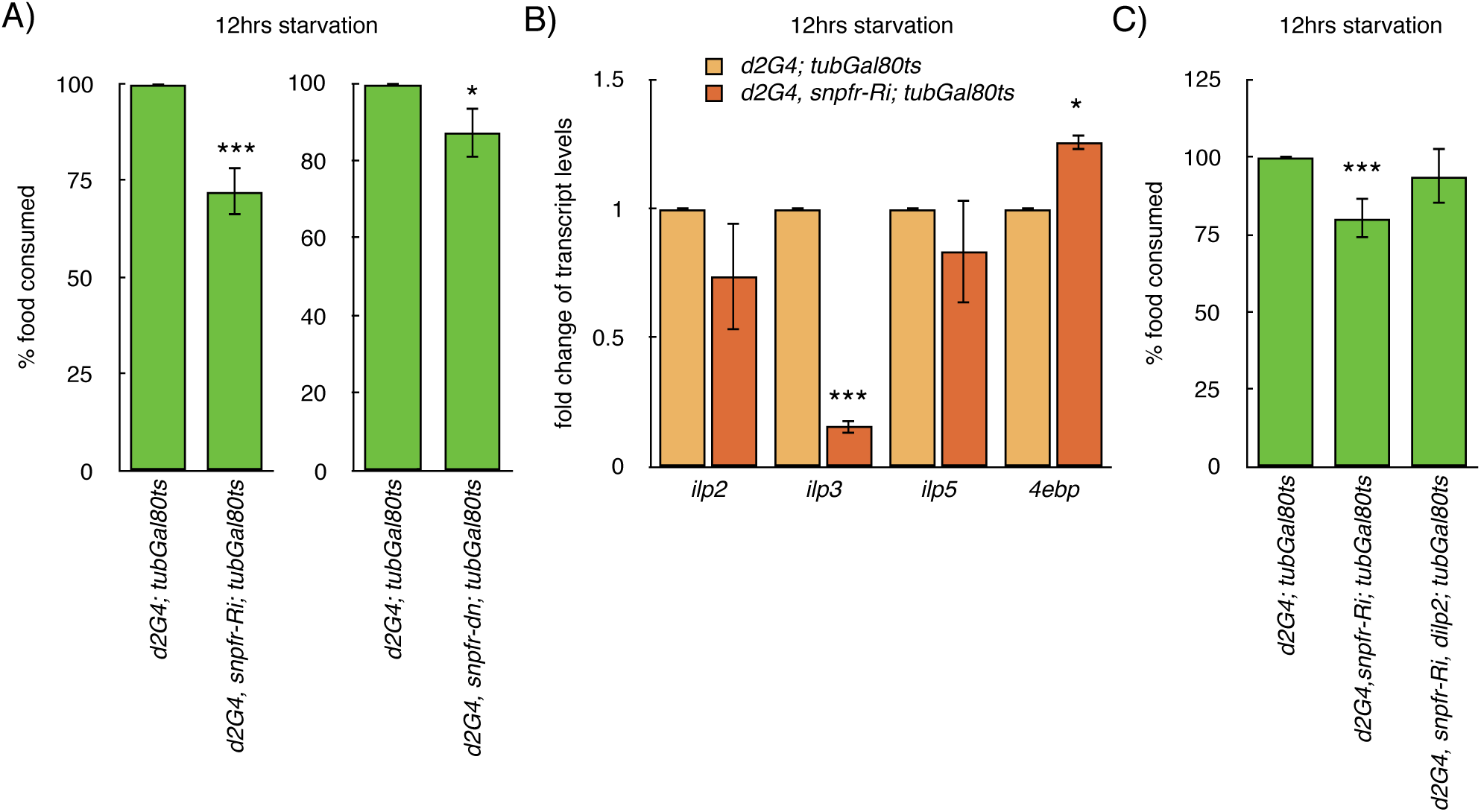
During early phases of food deprivation sNPF signalling in IPCs maintain insulin levels and enhance feeding. (**A**) blocking sNPF signalling in IPCs using *snpfr*-RNAi expression reduce feeding in response to 12 hours of starvation, whereas *snpfr*-RNAi expression in IPCs did not affect enhanced feeding in response to 36 hours of food withdrawal, expression of *UAS*-*snpfr*-*RNAi* was restricted using *tub*-*Gal80^ts^*, values show difference in feeding levels between control and *snpfr*-*RNAi* flies in percentage after indicated hours of starvation [n>18]. (**B**) changes in *dilp2*, *dilp3*, *dilp5* and *4e*-*bp* mRNA levels in response to blocking sNPF signalling in IPCs using *snpfr*-*RNAi* expression, values are fold change in transcript levels normalised to control genotype without *UAS*-*RNAi* [n=3]. (**C**) co-expressing *dilp2* in IPCs rescues *snpfr*-RNAi mediated reduction of feeding in response to 12 hours of starvation, expression of *UAS*-*snpfr*-*RNAi* and *UAS*-*dilp2* was controlled using *tub*-*Gal80^ts^*, values show difference in feeding levels between control, *snpfr*-*RNAi* and *snpfr*-*RNAi* + *dilp2* [n=14]. Expression of the various tranegenes were strictly controlled using *tub*-*Gal80^ts^*, the expression of the transgene was turned on 12 hours prior to the start of starvation by transferring flies from 18°C to 29°C. (P-value ^∗^<0.05; ^∗∗∗^ <0.005, n= number of biological replicates, error bars represents SEM)

As our results show that IPC function and maintenance of DILP levels are crucial for hunger-induced feeding at the initial phases of starvation, next, we checked if DILPs induce feeding by activating sNPF levels during early hours of food deprivation. We inactivated IPC function by expressing *reaper* just prior to starvation, this led to a reduction of *snpf* mRNA levels and starvation-induced feeding (Fig 6A and 3A). To confirm the role of insulin in maintaining orectic sNPF levels during early stages of starvation we blocked insulin receptor expression in the sNPF neurons, which showed a reduction of *snpf* mRNA levels and slight but significant reduction of hunger-induced food consumption (Fig 6B and C). However, at 36 hours of starvation blocking insulin signalling in sNPF neurons led to an increase in feeding, opposite to the effects seen during early stages of starvation (S6 Fig B). Blocking insulin signalling was shown to increase sNPF mRNA levels and feeding by a previous study (25), which is similar to the responses we observed by blocking insulin signalling in sNPF neurons during late stages of starvation. These findings confirm the role of IPCs in maintaining sNPF levels, crucial for food intake in response to hunger during early stages of starvation. Thus, in starved flies insulin and sNPF signalling acts in a feedback loop to enhance feeding responses in the initial stages. However, the feedback regulatory relationship between insulin and sNPF does not exist during later stages of starvation.

**Fig. 6.**
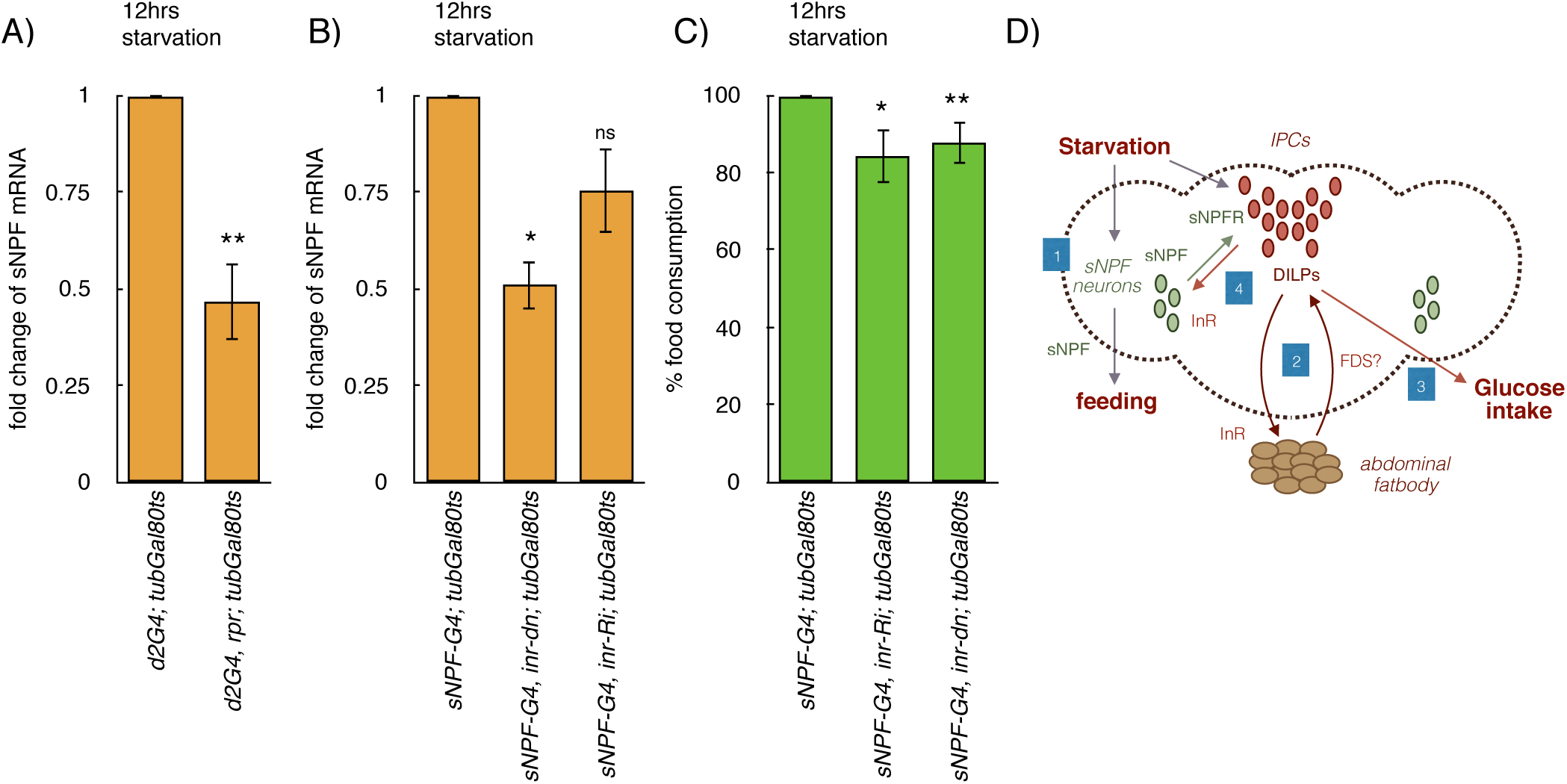
Insulin signalling in sNPF neurons aids in hunger induced feeding during early stages of starvation. (**A**) expression of *UAS*-*reaper* in IPCs led to a reduction of *snpf* mRNA levels, values show fold change in transcript levels compared to control genotype [n=3]. (**B**) blocking insulin signalling using *UAS*-*inr*-*RNAi* or *UAS*-*inr*-*DN* expression in sNPF expressing neurons led to reduction in *snpf* mRNA levels, values show fold change in transcript levels in comparison to control genotype [n=3]. (**C**) blocking insulin signalling using *UAS*-*inr*-*RNAi* or *UAS*-*inr*-*DN* expression in sNPF expressing neurons led to a reduction in hunger induced feeding responses in flies starved for 12 hours, values show % feeding levels [n>20 and n>5]. *P*-*value* ^∗^<*0.05*; ^∗∗^<*0.01*; ^∗∗∗^ <*0.005*, *n*= *number of biological replicates*, *error bars represents SEM).* (**D**) Model showing the regulation of feeding induced by food deprivation, and the crosstalk between IPCs and abdominal fatbody and the crosstalk between sNPF and IPC neurons. [1] Starvation acts on sNPF neurons, which through sNPF receptors on IPCs enhance DILP levels. [2] DILPs then act on abdominal fatbody to maintain DILP levels and feeding through a positive feed-back regulatory mechanism. Thus, DILP levels are maintained during early stages of starvation, despite the low nutrient levels. [3] High insulin signalling during the early phases of starvation aids in glucose uptake from circulation, however enhanced levels of anorexic insulin would affect feeding in response to starvation. [4] Further, DILPs act on sNPF neurons and maintain high sNPF levels and feeding during early stages of starvation, which aids in circumventing the anorexic effect of DILPs and induce feeding. The crosstalk between IPC neurons and abdominal fatbody maintains DILP levels and glucose uptake during early stages of starvation. And the crosstalk between sNPF and IPC neurons aids in feeding.

## Discussion

In complex multicellular organisms nutrient environment is sensed by neural, endocrine and enteric centers which control food intake and energy utilization for biological functions. In mammals, the hypothalamic arcuate nucleus (ARC) responds to changes in nutrient levels and control food consumption and energy expenditure (4, 36). Neural control of feeding is believed to be conserved in the animal kingdom as neural centers that regulate food intake and energy homeostasis, similar to the hypothalamic ARC, is present in most bilaterian animals (37). Though the control of feeding and maintenance of energy homeostasis by the brain is under intense investigation, the exact molecular mechanisms that function in the neural circuitry that controls feeding are not entirely understood yet.

### Insulin enhances feeding responses during nutrient deprivation

Insulin hormone suppresses food intake and is generally believed to act as a ‘satiety factor’ (38). However, in the neuron-specific *insulin receptor* knock-out mice (NIRKO) there were no significant changes to food intake in males (39). Earlier studies showed that the function of insulin as a satiety hormone is mediated in part by the hypothalamic AgRP (Agouti-related peptide)/NPY neurons (40). Despite this, mutant mice that lack InR in the AgRP/NPY neurons or the conventional *agrp*, *npy* double knockout mice did not show any defects to energy homeostasis and feeding capacity (41). However, ablation of AgRP and NPY neurons in the adult mice led to acute hypophagia and reduced body weight (42). Also, stimulation of AgRP neurons specifically in adult mice using optogenetic means triggered food intake (43). Thus, perturbation of the hypothalamic control of energy metabolism and food intake is subject to ‘developmental compensation’. The presence of such buffering mechanisms during development is understandable as control of metabolism and feeding are crucial to the survival of an organism and it’s reproductive fitness.

In Drosophila overexpression of *dilps* has been shown to block feeding in food deprived larvae showing that insulin acts as a ‘satiety hormone’ in flies as well (23, 25). Drosophila larvae show enhanced intake of less accessible or noxious food in starved conditions known as motivational feeding. Activation of insulin signalling reduced motivational feeding in starved larvae and blocking insulin signalling enhanced motivational feeding even in satiated larvae (2). Furthermore, enhanced insulin signalling in adult flies led to a decrease of *snpf* mRNA expression and reduced food intake (25). Thus, Drosophila insulin has been shown to act as a negative regulator of both hunger induced and baseline food intake.

Contrary to the belief that insulin acts as a ‘satiety hormone’, we show that during initial phases of food withdrawal insulin levels are maintained and trigger feeding responses in Drosophila (Fig 3). We have modulated insulin signalling specifically in the adult animals to avoid developmental perturbations and possible compensatory effects, which may have revealed functions of insulin unknown to us before. However, it may be noted that comparable results as published earlier were seen by enhancing or blocking insulin signalling right from the embryonic stages (data not shown) (23, 25). Insulin was found to act as an orectic hormone only during early stages of starvation and during late starvation insulin acts as an anorexic hormone as reported earlier (S2 Fig B). However, the mechanisms that sustain insulin levels during early phases of starvation should be strong enough to overcome opposing effects of nutrient deprivation on insulin levels. As discussed in the next section, multiple feedback signalling loops involving sNPF neurons and abdominal fatbody would sustain a critical level of insulin signalling despite the negative energy status of the organism.

### Multiple feedback loops that maintain feeding

Feedback loops act in various cell signalling pathways to switch on or switch off gene expression efficiently. For example, positive feedback loops can lead to timely developmental decisions called developmental switches (44-46). In physiological and metabolic contexts also feedback regulation helps in mounting immediate responses to fluctuations in various environmental factors (47-51). Insulin signalling also acts through various feedback signalling mechanisms to manage outcomes in different physiological and developmental contexts (52). Insulin receptor levels are transcriptionally regulated by insulin signalling through the transcription factor FOXO (30, 31). In addition, insulin is known to exert feedback regulation on Insulin Receptor Substrate (IRS), AKT, Target of Rapamycin (TOR) and Hypoxia-inducible factor-1α (Hif1α), which in turn regulates the activity of insulin signalling pathway (53, 54). Here in this paper, we unravel multiple feedback signalling loops activated by insulin in various tissues that manage nutrient homeostasis, which together enhances feeding in response to hunger.

### Feedback regulatory mechanism that involves IPCs and abdominal fatbody that aids in feeding during early stages of starvation

Our results show that insulin signalling in the abdominal fatbody aids in feeding during early stages of starvation, by maintaining IPC specific *dilp* gene expression and insulin signalling. Fat derived signals like Upd2, Eiger, CCHamide and DILP6 has been shown by previous studies to influence IPC function and DILP levels based on the growth and nutrient status of Drosophila (18, 19, 26, 27, 34). Though loss of *upd2* led to the accumulation of DILP2 in the IPCs and resulted in growth phenotypes no effect on feeding has been reported thus far (18). Other fat derived signals DILP6 and Eiger are expected to increase in response to 16-18 hours of food deprivation (19). We predicted that these humoral signals may be meaningful in regulating feeding during early starvation. However, manipulation of the levels of *upd2*, *dilp6* and *eiger* gene expression did not show any impact on feeding during early starvation, hence a yet unknown fat derived signal or multiple fat derived signals regulate starvation induced feeding. Thus, through a feedback regulatory mechanism insulin acts on abdominal fatbody to maintain *dilp* mRNA levels and hunger driven feeding during early stages of starvation (Fig 4, Fig 6D).

### sNPF and DILP signalling act in a positive feedback loop to enhance feeding during early stages of starvation

Nutrient deficiency triggers hunger-induced feeding responses in flies, similar to vertebrates. In Drosophila, the loss of snpf gene function led to the reduction of body size and food intake (35). sNPF stimulates dilp gene expression in the larval IPCs that leads to growth in flies (20). Despite the fact that sNPF activates feeding and its action on IPCs trigger insulin gene expression during growth, the involvement of IPCs in the sNPF mediated feeding is still not clear.

Our results establish that sNPF neurons and the IPCs are involved in a positive feedback loop, where sNPF activates IPCs, and IPCs in turn would activate *snpf* gene expression and feeding during the initial phases of starvation (Fig 6D). Previous studies show that insulin signalling has an inverse regulatory relationship with the nutritional status of an organism (55). However, our experiments show that during initial phases of starvation sNPF acts on IPCs to maintain *dilp* gene expression, insulin signalling and enhance the drive to feed (Fig 5A and B). This response is similar to the effects of sNPF signalling in IPCs during larval growth (20). Unlike early stages of starvation sNPF signalling in the IPCs during later stages had no effects on feeding (S6 Figure A). Furthermore, we show for the first time that IPC function in sNPF neurons is inevitable for *snpf* mRNA expression and food intake during the early hours of starvation (Fig 6A and B). Blocking insulin signalling led to an increase in feeding in flies subjected to food withdrawal for 36 hours consistent with the previous reports (23, 25). Thus, sNPF and IPC neurons are involved in a positive feedback loop during the initial phases of starvation where sNPF activates IPCs, which in turn maintain *snpf* gene expression and feeding (Fig 6D). Here again, deviation from the expected results could be attributed to the circumvention of developmental redundancy and limiting possible non-physiological effects.

Expression of *snpf* gene expression is suppressed by DILPs in satiated conditions (25). Also, chromatin immunoprecipitation experiments show that FOXO interacts with *snpf* promoter during food deprivation. However, during early stages of starvation insulin signalling may not be as low as expected to promote the binding of transcription factor FOXO to sNPF promoter [this study and (25)]. Our results show that insulin signalling in the sNPF neurons, through a hitherto unknown mechanism, aid in gene expression of sNPF which stimulates feeding during early stages of starvation (Fig 6A and B). However, during late stages of starvation low levels of insulin may induce sNPF gene expression, most probably through a FOXO dependent mechanism as proposed previously.

### Why is insulin levels maintained high during starvation?

We made a perplexing but interesting observation that adult fruit flies depend on an anorexic hormone insulin to elicit hunger induced feeding responses. Maintaining sNPF levels alone should suffice to induce feeding in response to starvation and the simultaneous maintenance of high levels of insulin is counterintuitive to the known functions of insulin. Based on our results, we propose that maintaining insulin levels during the early stages of starvation has two physiological roles: i) to help in the utilization of energy sources to manage normal activities during early starvation and ii) to trigger a rapid feeding response during early stages of starvation. Previous studies have shown that during early stages of starvation the drive to feed is very high, which would trigger behavioral responses like foraging and motivated feeding (2). The drive to feed in response to hunger must have a quick onset and should be sustained until new food sources are found, which would help from circumventing the ill-effects of long-term starvation. Despite the known anorectic functions insulin signalling is crucial for driving *snpf* gene expression and feeding during early stages of starvation (Fig 6). The regulatory feedback loop which involves sNPF neurons and the IPCs would aid in maintaining high sNPF levels and trigger a rapid onset of feeding during early stages of starvation. The hunger inducing activity of insulin is not entirely surprising as enhanced insulin levels is known for a long time to induce feeding during physiological and disease states (56-58). In short, we identify an ‘orexigenic’ phase of activity of insulin during early stages of starvation.

## Materials and Methods

### Fly Strains

Fly stocks were maintained in vials at 25°C with 12h:12h light:dark cycle. The flies with *tub* - *Gal80^ts^* were maintained at 18°C to express active Gal80 to restrict the expression of Gal4, these flies were shifted to 29°C only 12 hours prior to starvation and during starvation (for details check S1 Figure). The standard fly food consisted of cornmeal, dextrose, yeast, agar and Nipagen. Mifepristone or RU486 (Sigma) was added (0.5mg/ml) whenever Gene-Switch Gal4 experiments were performed. For Gene-Switch Gal4 experiments fly stocks were maintained in vials at 25°C with standard fly food, these flies were shifted to Mifepristone vials only 12 hours prior to starvation and during starvation (for details check S3 Figure). Following fly strains were obtained from Bloomington Drosophila Stock Center - *UAS*-*InRdn* (8251), *inr^E19^/TM2* (9646), *inr^GC25^/TM6* (9554), *S_1_32*-Ga14 (8527) *S_1_106*-Ga14 (8151), UAS-*upd2 TRiP* lines (33988 and 33949). The RNAi lines *UAS*-*inr*-*Ri* (v992), UAS-*dilp6*-*Ri* (v102465) UAS-*eiger*-*Ri* (v45253, v108814) were obtained from VDRC. *dilp2*-Ga14, UAS-*mCD8GFP/CyO*, *tub*-Ga180*^ts^/TM2*, *arm*-Ga14, UAS-*dilp2*, UAS-*TrpA1*, UAS-*rpr*, UAS-*PTEN*, UAS-*Dilp6*, UAS-*sNPFR*-*Ri*, UAS-*Kir2.1*, UAS-*PH*-*GFP* and *w^1118^* were obtained from Steve Cohen’s lab. UAS-*snpfr*-*DN* was a kind gift from Kweon Yu. *sNPF*-Gal4 was a gift from Michael Rosbash.

### Adult Feeding Assays

Flies containing *tub*-*Gal80^ts^* were reared at 18°C, 5 day old male flies were shifted to 29°C incubator 12 hours prior to the experiment and retained at 29°C till the end of the experiment. The flies were then kept for 12 or 24 hours in either fed conditions using normal food vials containing yeast paste or starved conditions in 1% agar vials. Flies were then allowed to feed on yeast paste with Orange G (Sigma) for 30 minutes. The amount of food consumed were analysed by lysing equal number of flies per tube using Bullet Blender Storm (Next Advance, NY). The flies were homogenized using zirconium oxide beads and homogenization buffer (0.05% Tween 20), the supernatant was used for the colorimetric estimation at 478nm using TECAN Infinite M200 pro multi-mode plate reader in 96 well plates. The absorbance was plotted normalized against the control.

### Immunostaining, Confocal Microscopy and image analysis

DILP2 polyclonal antibody was raised using a peptide corresponding to amino acids 108–118 of the DILP2 sequence (TRQRQGIVERC) as an immunogen in rabbits (Eurogentec, Belgium) (17). For whole mount immunostaining, fatbody from 5-6 day old male flies were dissected in Shields and Sang M3 Insect medium (Sigma), and the brains were dissected using ice-cold PBS. Samples were then fixed in 4% paraformaldehyde (PFA) (Sigma) for 20 minutes at RT. Primary antibodies anti-GFP (Invitrogen) or anti-DILP2 were added at a dilution of 1:500 was added and incubated overnight at 4°C. Samples were then incubated with secondary antibody (Alexa Fluor^®^ 488 Goat Anti-Rabbit IgG Invitrogen) for not more than 2 hours at RT. Samples were then mounted in mounting medium (SlowFade^®^ Gold Antifade Reagent with DAPI, Invitrogen). Imaging was done using Leica DM6000B upright microscope. ImageJ software was used to process and analyse the images. The confocal Z stack of IPCs each with 2 μm step size and same laser power settings were used across samples to measure the intensity of florescence.

### Quantitative real-time PCR

Fly heads or body samples were flash frozen in liquid nitrogen. Total RNA was isolated using RNeasy Plus mini kit (Qiagen). RNA samples were reverse transcribed using SuperScript III (Invitrogen), generated cDNA was used for real time RT-PCR using Bio-Rad CFX96^TM^. Primers for *rp49* and *actin* were used to as reference genes to quantify the target amplification. A minimum of three independent biological samples were analysed for each condition, and qPCR plates were setup using 3 technical replicates from each biological replicate.

List of primers used for qPCR

*dilp2 FP*: *GGCCAGCTCCACAGTGAAGT*
*dilp2 RP*: *TCGCTGTCGGCACCGGGCAT*
*dilp3 FP*: *CCAGGCCACCATGAAGTTGT*
*dilp3 RP*: *TTGAAGTTCACGGGGTCCAA*
*dilp5 FP*: *TCCGCCCAGGCCGCAAACTC*
*dilp5 RP*: *TAATCGAATAGGCCCAAGGT*
*rp49 FP*: *GCTAAGCTGTCGCACAAA*
*rp49 RP*: *TCCGGTGGGCAGCATGTG*
*act5C FP*: *CACACCAAATCTTACAAAATGTGT*
*act5C RP*: *AATCCGGCCTTGCACATG*
*4ebp FP*: *CACTCCTGGAGGCACCA*
*4ebp RP*: *GAGTTCCCCTCAGCAAGCAA*
*inr FP*: *AACAGTGGCGGATTCGGTT*
*inr RP*: *TACTCGGAGCATTGGAGGCAT*
*snpf FP*: *CCCGAAAACTTTTAGACTCA*
*snpf RP*: *TTTTCAAACATTTCCATCG*

### Western blotting

Anti-Akt, anti-phospho-Akt (Ser505) and anti-actin antibodies were purchased from Cell Signalling Technology. Adult whole flies (5 per condition) was homogenised in RIPA buffer with Protease Inhibitor cocktail (Sigma). The homogenate was centrifuged and the supernatant was added with loading buffer containing SDS for 10 minutes at 95°C. The denatured lysates were run on 10% SDS-PAGE, transferred to nitrocellulose membrane, incubated with following primary antibodies: phospho- Akt (Ser505) and total Akt diluted 1:1000 in 5% BSA in TBST and anti-actin antibody was diluted 1:1000 in 5% non fat dry milk in TBST, were incubated in 4°C overnight. Following the washes the blots were treated with HRP conjugated secondary antibodies and the immunoblotting signals were visualised using Immobilion Western Chemiluminescent HRP Substrate (Merck, Millipore). Bands were visualised using the Bio-Rad ChemiDoc^™^ XRS+ imaging system.

### Statistics

All the feeding behaviour experiments were run along with the control genotypes in parallel, with a minimum of n=06 per genotype, each n is a set of 7 flies, and the data is the summation of multiple independent experiments. Data is represented as a mean ± SEM and Student’s *t*-test (two tailed, two sample unequal) was used to compare the statistical significance between the control and the test group. Feeding assays and q-PCR experiments were analysed using the above mentioned tests.

## Acknowledgments

We thank Steve Cohen, Michael Rosbash, Kweon Yu, Tina Mukherjee, VDRC and Bloomington Stock Center for flies. We are thankful to Nisha Kannan, Smitha Vishnu, Himani Pathak Niyas Rehman and Jervis Fernandes for critical comments on the manuscript. SRS is a senior research fellow of DST-India Inspire PhD program and JV is supported by Ramanujan Fellowship from DST-SERB, India. We thank IISER TVM, MHRD, Government of India for the generous intramural financial and infrastructural support.

